# Heat Stress Induces Locus-Specific DNA Hypomethylation Linked to Immune Regulation in Lactating Holstein Cows

**DOI:** 10.64898/2026.03.23.713208

**Authors:** Gabriel Costa Monteiro Moreira, Alexis Ruiz González, Mélodie Joigner, Valentin Costes, Aurélie Chaulot-Talmon, Francesca Ali, Lorraine Bourgeois-Brunel, Hélène Jammes, Daniel E. Rico

**Author notes:** Corresponding autors: Gabriel Costa Monteiro Moreira, Daniel E. Rico.

## Abstract

Epigenetics may play a crucial role in livestock adaptation to environmental challenges like heat stress. In recent years, a growing number of studies have investigated the epigenetic mechanisms underlying dairy cow adaptation to heat stress. However, there is still limited knowledge about the effects of heat stress on immune cells and immune-related phenotypes. Herein we aim to identify heat-stress induced DNA methylation variations on blood methylome potentially affecting regulatory regions and associated phenotypes. Blood samples were collected and peripheral blood mononuclear cell (PBMC) isolated from four cows before (D0) and after (D14) a 14-d heat stress challenge (cyclical THI 72-82) and, from four cows kept in thermoneutral conditions (THI 61-64). Heat-stressed cows had *ad libitum* access to diets supplemented with adequate levels of vitamin D₃ and Ca (12,000 IU/kg of vitamin D₃ and 0.73% Ca, respectively). To eliminate confounding effects due to differences in nutrient intake, cows maintained under thermoneutral conditions were pair-fed (PF) to their heat-stressed counterparts and received adequate concentrations of vitamin D₃ and Ca as well. Reduced representation bisulphite sequencing (RRBS) was used to profile PBMC’s methylome. Differential methylation analysis was performed using *methylKit* and *DSS* software’s (Δmeth ≥ 25%, adjusted p-value < 0.01), retaining only commonly detected differentially methylated cytosines (DMCs). A total of 2,908 DMCs were identified when comparing pre- and post-heat stress samples. After excluding 649 DMCs that were also detected under thermoneutral conditions, as these changes were likely associated with feed restriction inherent to the pair-feeding design rather than with heat stress per se, 2,259 heat stress-specific DMCs remained, predominantly hypomethylated. About half of the DMCs are annotated in intronic and intergenic regions; known to harbor regulatory elements. By intersecting the DMRs with publicly available functional annotation data, we observed hypomethylation on regulatory regions putatively affecting cow’s immune system. As an example, we identified a loss of methylation within an enhancer region of the *MSN* gene, which is involved in lymphocyte homeostasis, and a loss of methylation in the promoter region of *MECP2*, a well-established epigenetic regulator with a central role in chromatin organization and gene expression. These findings highlight the impact of heat stress on dairy cow immunity and provide new insights into its epigenetic regulation under environmental stress.

**Interpretative summary:** This study examined DNA methylation changes induced by heat stress in dairy cows to elucidate epigenetic mechanisms of thermal adaptation. Using RRBS on PBMCs, 2,259 heat stress-specific differentially methylated cytosines were identified, predominantly hypomethylated and enriched in regulatory regions. Functional annotation highlighted immune-related pathways, including hypomethylated regulatory regions near genes (e.g., *MSN, ZBTB33, SLC25A5, GNAS, FAM3A,* and *MECP2*) associated with immune function. These findings indicate that heat stress induces targeted epigenetic modifications potentially affecting immune regulation in dairy cows.

## Introduction

Climate change is increasing the frequency, intensity, and duration of heat load events, placing unprecedented stress on livestock production systems and threatening the sustainability of dairy cattle operations worldwide (Nardone et al., 2010). High ambient temperatures and humidity often quantified using the temperature-humidity index (THI), compromise thermoregulation in high-producing dairy cows, resulting in elevated core body temperature, increased respiration rates, and behavioral adaptations that are insufficient to fully dissipate thermal load, resulting in heat stress (Chapman et al., 2017).

Heat stress is well recognized for its detrimental effects on dairy cow well-being and productivity, contributing to reductions in dry matter intake, milk yield and milk solids, which are associated with altered nutrient partitioning, as well as increased intestinal permeability and systemic inflammation (Wheelock et al., 2010; Ruiz-González et al., 2023). At the organ and cellular level, heat stress alters blood flow distribution, leading to intestinal vasoconstriction, intestinal hypoxia, and compromised tight junction integrity (Lambert, 2009). This impairment of barrier function increases the passage of luminal contents such as bacterial lipopolysaccharide (LPS) into the bloodstream (i.e., endotoxemia), provoking systemic immune activation and inflammation (Koch et al., 2019; Ruiz-González et al., 2023). Endotoxemia and inflammation can significantly influence nutrient partitioning and endocrine function, increasing insulin secretion despite lower feed intake, shifting glucose use towards immune response (Ganeshan and Chawla, 2014) rather than milk production (Kvidera et al., 2017). Similarly, we and others have shown hyperinsulinemia, increased gut leakiness, and systemic inflammation during heat stress are associated with reduced productivity during heat stress (Fontoura et al., 2022; Ruiz-González et al., 2023).

In addition to these acute physiological consequences, heat stress has persistent effects on animal performance that may be mediated by long-lasting changes in gene regulation. One well-studied model of this phenomenon is *in utero* heat stress, where late-gestation exposure of pregnant dams to high ambient THI results in profound, lifelong deficits in offspring growth, immunity, reproduction, productivity and markers of longevity (Monteiro et al., 2016; Skibiel et al., 2018; Laporta et al., 2020; Kipp et al., 2021). Importantly, as determined by pair feeding studies, some of these effects (e.g., calf growth and immunity) are not solely explained by reduced feed intake of the dams, but are indeed the specific result of hyperthermia (Almoosavi et al., 2020). (Monteiro et al., 2016) demonstrated that heifers exposed *in utero* to late-gestation maternal heat stress were smaller and yielded less milk and milk components during their first lactation compared with heifers from cooled dams, which may be partially explained by the well-documented alterations the morphology of the mammary gland (Skibiel et al., 2018).(Laporta et al., 2020) showed that relative to their cooled counterparts, heat stressed dry cows birthed offspring with significantly reduced milk yields in the first and subsequent lactations, as well as reduced productive lifespan. Importantly, recent work from (Boucher et al., 2025) reported persistent methylation changes in whole blood of *in utero* heat-stressed dairy calves from birth to weaning, suggesting possible epigenetic mechanisms which could explain at least in part the observed performance losses of these animals.

The body of work on *in utero* heat stress highlights the potential for environmental heat load to exert lasting effects via epigenetic mechanisms, yet our understanding of how heat stress experienced during adult life influences epigenetic regulation in dairy cows remains limited. Prior research has identified heat-induced differences in peripheral blood mononuclear cells (PBMCs) phenotypes, likely in response to heat stress-induced endotoxemia (Koch et al., 2025). However, to date, these studies have not systematically evaluated methylation changes in lactating cows exposed to controlled experimental heat stress compared with thermoneutral conditions using a per-feeding strategy, nor have they advanced in linking these changes to concurrent immune activation and metabolic dysregulation through integration of functional annotation data.

Understanding adult heat stress epigenomics is particularly relevant given the pronounced immune activation and metabolic adjustments that characterize the heat-stressed phenotype. Immune activation, whether driven by endotoxin translocation or direct thermal effects, is energetically expensive, altering nutrient allocation and production performance. Moreover, immune and metabolic pathways are tightly regulated at the transcriptional level, and epigenetic modifications such as DNA methylation are known to modulate gene expression in response to environmental stimuli (Halli et al., 2025). Therefore, characterization of methylation profiles in key immune cells during heat stress exposure may illuminate regulatory mechanisms linking environmental challenges to cellular function.

Recent studies report heat stress-associated differentially methylated regions (DMRs) and gene-specific methylation shifts, suggesting that targeted methylation loci, or small panels of CpGs, could serve as robust biomarkers to identify animals that experienced clinically relevant heat stress (Del Corvo et al., 2021; Halli et al., 2025). Thus, the objective of the present study was to evaluate the impact of experimentally induced moderate to severe heat stress on DNA methylation profiles in PBMCs of lactating dairy cows. We hypothesized that marked alterations in methylation patterns would occur under heat stress, potentially leading to changes in immune activation and cellular stress responses, and that these methylation changes would provide insight into mechanisms underlying heat stress-induced immune and metabolic phenotypes.

## Material and Methods

### Animal care conditions, management, phenotypes measurements and blood sample collection

Animal care conditions and management practices have been previously published (Ruiz-González et al., 2023). Briefly, multiparous Holstein cows were splitted into two different groups, balanced for parity, days in milk (DIM), and milk yield. The experiment was conducted between October and December 2020 at the CRSAD research farm (Deschambault, QC, Canada), where cows were housed in a tie-stall barn, in a climate-controlled chamber with a 12-h light and dark cycle. Four heat-stressed (HS) cows experienced a cyclical daily temperature-humidity index (THI) calculated according to (Schüller et al., 2014) varying between 72.0 and 82.0 via automated control of chamber temperatures (Edgetech Instrument Inc.). Temperature control in the heat challenge was as follow: gradual increase from 29 to 39 °C between 0800 and 1400 h at a rate of 1.6 °C per hour, followed by a constant temperature phase from 1400 to 1700 h. The temperature was then progressively reduced back to 29 °C between 1700 and 2300 h at the same rate, and maintained at 29 °C from 2300 to 0800 h. The relative humidity was not controlled and ranged from 20 to 50%. This daily temperature cycle was applied throughout the 14-day experimental period. Four thermoneutral cows (TN) experienced thermoneutral conditions with temperature maintained at 20°C and the relative humidity ranging from 55 to 64% (temperature-humidity index = 61.0 to 64.0) (Ruiz-González et al., 2023).

To eliminate confounding effects due to differences in nutrient intake, heat-stressed cows had *ad libitum* access to diets supplemented with adequate levels of vitamin D₃ and Ca (12,000 IU/kg of vitamin D₃ and 0.73% Ca, respectively while cows maintained under thermoneutral conditions were pair-fed (PF) to their heat-stressed counterparts and also received adequate concentrations of vitamin D₃ and Ca. Both group of cows were milked twice daily (0700 and 1700 h) in their respective stalls. Morning and afternoon milk samples were pooled by cow; milk yield was determined at D0 (day 0; baseline - immediately before the challenge) and D14 (day 14, end of the challenge) using an integrated milk meter (Flomaster Pro; DeLaval). An additional aliquot was stored in 4°C freezer with preservative (2-bromo-2-nitropropane-1,3-diol) until analyzed for milk components. A detailed description of the animal cohort, experimental conditions and phenotypic measurements is available in (Ruiz-González et al., 2023).

Animal performance, as well as metabolic and inflammatory markers, were assessed in heat-stressed (HS) and thermoneutral (TN) cows, as previously described (Ruiz-Gonzalez et al., 2023). Both groups were monitored for rectal temperature, respiratory rate, dry matter intake (DMI), and milk yield over a 14-day period.

Blood samples were collected from the coccygeal vein at D0 and D14. Samples were collected into disposable glass culture tubes containing 250 U of sodium heparin, immediately placed on ice, and centrifuged within 20 min at 1,500 × g at 4°C for 20 min. Extracted peripheral blood mononuclear cells (PBMCs) were aliquoted and frozen at −20°C. PBMCs aliquots were shipped to INRAE Jouy-en-Josas (BREED unit) headquarters for genomic DNA extraction.

### Genomic DNA extraction, library preparation and sequencing

Genomic DNA (gDNA) was extracted from PBMCs following the standard proteinase K protocol. After extraction, DNA integrity was evaluated on agarose gel (1%) electrophoresis for all samples and, DNA purity and concentration were assessed on Nanodrop (Thermo Fisher Scientific) and QuBit fluorometer (Thermo Fisher Scientific), respectively.

For this study, sixteen reduced representation bisulfite sequencing (RRBS) libraries were prepared: four at D0 and four at D14; before and after heat stress challenge, and four at D0 and four at D14; at the same timepoint from cows under thermoneutrality conditions. As described in (Costes et al., 2022), genomic DNA was cleaved by MspI enzyme at CCGG sites, generating fragments with CG dinucleotides at both ends followed by end repair step and Illumina adaptors ligation. The fragments were then selected by their size keeping DNA fragments with a size between 150 - 400 bp (40–290 bp genomic DNA fragments + adapters), using SPRIselect magnetic beads (Beckman-Coulter, Villepinte, France). After size selection, bisulfite conversion was performed using the EpiTect bisulfite kit (Qiagen, Les Ulis, France) following manufacturers’ instructions and, amplified with Pfu Turbo Cx hotstart DNA polymerase (Agilent, Les Ulis, France) using 14 PCR cycles and Illumina index. PCR products were analyzed using the Agilent TapeStation system (Agilent Technologies, Santa Clara, CA, USA) to assess library quality and homogeneity. Paired-end (PE) sequencing with a read length of 100 bp, was performed in Illumina NovaSeq 6000 (Integragen-OncoDNA, Evry, France), aiming a minimum of 30M PE reads by library.

### Data analysis

Quality control of sequencing data, alignment, CpG detection and filtering, methylation estimation and differential methylation analysis were performed as described in (Costes et al., 2022). Briefly, after sequencing, quality control on raw reads was performed using FastQC (https://www.bioinformatics.babraham.ac.uk/projects/fastqc/) and then, reads with a Phred score ≤ 20, shorter than 20 nucleotides and adaptor sequences were trimmed using Trim Galore! (v0.4.4; https://www.bioinformatics.babraham.ac.uk/projects/trim_galore). Bisulfite conversion rate was estimated from unmethylated cytosine added *in vitro* during the end-repair step. Quality-trimmed reads were aligned against the bovine reference genome (ARS-UCD1.3 - GCA_002263795.3) using Bismark(Krueger and Andrews, 2011) v0.20.0 (default parameters) with Bowtie 1.2.1(Langmead et al., 2009). Only CpGs covered by a minimum of 10 and maximum of 500 uniquely mapped reads (CpGs_10-500_) and commonly detected in all the libraries (*common* CpGs_10-500_) were retained for subsequent analysis. Additionally, *common* CpGs_10-500_ that co-localizes with a putative sequence polymorphism affecting either the C and/or G were removed to avoid confounding effects (SNP panel described in (Daetwyler et al., 2014). For each remaining *common* CpG_10-500_ was assigned a methylation percentage per sample calculated using the “*methylation_extractor*” function from Bismark which calculates methylation level by counting methylated (C) and unmethylated (T). Non-supervised hierarchical clustering was performed using the FactoMineR package(FactoMineR: Multivariate Exploratory Data Analysis and Data Mining, 2006) on the common CpGs10-500 methylation matrix.

Differentially methylated cytosines (DMCs) were identified using two independent algorithms, methylKit (v1.0.0) and DSS (v2.14.0), both run with default settings. For methylKit, a CpG site (among the common CpGs_10-500_) was considered a DMC if it showed an average methylation difference of at least 25% between the two experimental groups and an adjusted p-value (q-value) below 0.01, using SLIM correction. For DSS, a CpG site was considered differentially methylated if it also exhibited a minimum methylation difference of 25%, and the adjusted p-value below 0.01 - calculated using the Independent Hypothesis Weighting (IHW) method (in this case, the alpha parameter was set to 5%, and the average methylation level per group was used as a covariate).

To maximize the detection of methylation changes putatively induced by heat stress, different filtering strategies were applied. In the heat stress group, only CpG sites (CpGs_10-500_) that were identified as DMCs by both algorithms were retained. In contrast, for the thermoneutral group, all CpGs_10-500_ identified as DMCs by either algorithm (methylKit or DSS) were retained. This combined set from the thermoneutral group was then used to filter out overlapping DMCs from the heat stress group, thereby enriching for CpG sites likely associated with the heat stress response.

### Genomic features, regulatory elements, gene ontology and pathway analysis

All *common* CpGs_10-500_ as well as the DMCs were annotated relative to gene features, CpG density and repetitive elements as described in (Costes et al., 2022) and (Perrier et al., 2020). The reference files utilized were downloaded from Ensembl (ftp://ftp.ensembl.org/pub; release 100). Briefly, DMCs were assigned in transcription start site (TSS) if they are located − 100 to + 100 bp relative to the transcription start site (TSS); in promoter if located −2000 to − 100 bp relative to the TSS; in transcription termination site (TTS) if located – 100 to + 100 bp relative to the TTS; in shore if located up to 2000 bp from a CGI, and in shelf, if located up to 2000 bp from a shore. A site/region was considered to belong to a CGI (respective shore and shelf) if an overlap of at least 75% was observed between the site/fragment and the CGI (respective shore and shelf). A site was considered as being overlapped by a repetitive element whatever the extent of this overlapping.

The list of genes targeted by the final set of DMCs (after all filtering steps) underwent functional enrichment analysis using the ShinyGO v0.82 web tool, with *Bos taurus* selected as the reference genome. As the background, we provided the list of genes targeted by all common CpGs_10-500_, thereby accounting for the actual genomic space assessed in our study. Enrichment was conducted using the STRING database (within ShinyGO v0.82 web tool), focusing on protein-protein interaction networks and associated functional annotations. Only terms or pathways involving at least two genes and with an FDR < 0.05 were considered significant. To further support the biological relevance of the enriched terms and prioritized candidate genes, we carried out an extensive search of the scientific literature and available genomic databases.

To gain deeper insight into the functional role of the identified markers, we grouped the DMCs into regions sharing the same methylation status. Differentially methylated regions (DMRs) were identified and defined as regions containing at least three DMCs with similar methylation status (hypo- or hypermethylated) with an inter-distance between each DMC of 100 bp or less. We then intersected these regions and our list of pre-selected candidate genes with regulatory features from Ensembl (Ensembl Regulatory features – ARS-UCD1.3; https://ftp.ensembl.org/pub/release-113/regulation/bos_taurus/ARS-UCD1.3/annotation/Bos_taurus.ARS-UCD1.3.regulatory_features.v113.gff3.gz).

## Results

### Effects of Heat Stress on Physiological and Productive Traits

The impact of the heat stress on animal performance, and metabolic and inflammatory markers has already been published (Ruiz-González et al., 2023). Briefly, none of the variables evaluated was significantly different between the HS and TN groups on D0. Rectal temperatures and respiratory rates registered on D14 at 5pm were 4 and 240% higher, respectively, in HS compared with TN cows (all *P* < 0.01). Dry matter intake (DMI) decreased progressively in both the HS and the TN groups, reaching a nadir on D7 (i.e., −33%; Time *P* < 0.001). Milk yield was also reduced progressively over time, but despite similar levels of DMI, milk yield was on average 14% lower in HS cows from d 3 to 14 (*P* < 0.001). The plasma concentrations of insulin, lipopolysaccharide-binding protein, tumor necrosis factor alpha, C-reactive protein and of fecal calprotectin, were 46, 96, 202, 201, and 27% higher, respectively, in HS relative to TN cows (all *P* < 0.01).

### DNA methylation profiles unravel marked epigenetic effects of heat stress

Sequencing metrics for all libraries are detailed in Supplementary File S1. As described in the Methods section, the bisulfite conversion rate was estimated using unmethylated cytosines introduced *in vitro* during the end-repair step, yielding a high average conversion efficiency of 99.9%. On average, 48 million paired-end reads were generated per library, with an overall mapping rate of 86.03%. Of these, 32.21% represented uniquely mapped reads, consistent with previous studies (Del Corvo et al., 2021; Costes et al., 2022). This relatively low proportion of uniquely mapped reads is expected, given that RRBS preferentially targets CpG-rich regions, including repetitive elements, and that approximately half of the bovine genome consists of repeat sequences (Adelson et al., 2009; Elsik et al., 2009), which increases the likelihood of multi-mapping.

Despite this, uniquely mapped reads aligned with an average of 3.0 million CpG sites per library (from ∼28 million in the bovine genome), of which 71% were covered by between 10 and 500 reads (CpGs_10-500_) and retained for downstream analyses. The global average DNA methylation level in PBMCs was estimated at 54%± 1.79% (Supplementary File S1).

We used the set of CpGs_10-500_ consistently detected across all libraries (see Methods) prior to filtering to estimate the relative proportions of major immune cell types through a reference-based deconvolution approach. Specifically, we applied the *EpiDISH* R package (Teschendorff et al., 2017), which implements robust algorithms for cell-type deconvolution, in combination with a custom reference panel of DNA methylation profiles. Our reference panel was generated from purified immune cell populations (lymphocytes, monocytes, and neutrophils) isolated from blood samples of Holstein cows (Costes et al., unpublished data; manuscript in preparation). By applying this approach, we did not observe significant differences (t-test; *p* > 0.05) in the proportions of the three immune cell types before and after the heat stress challenge (Figure 1). Although not statistically significant, in the heat stress group we detected a slight decrease in the proportion of lymphocytes accompanied by a slight increase in neutrophils (Figure 1A, B, C). In contrast, this trend was not observed in the thermoneutral group, except for a slight reduction in the proportion of monocytes (Figure 1D, E, F).

**Figure 1.**
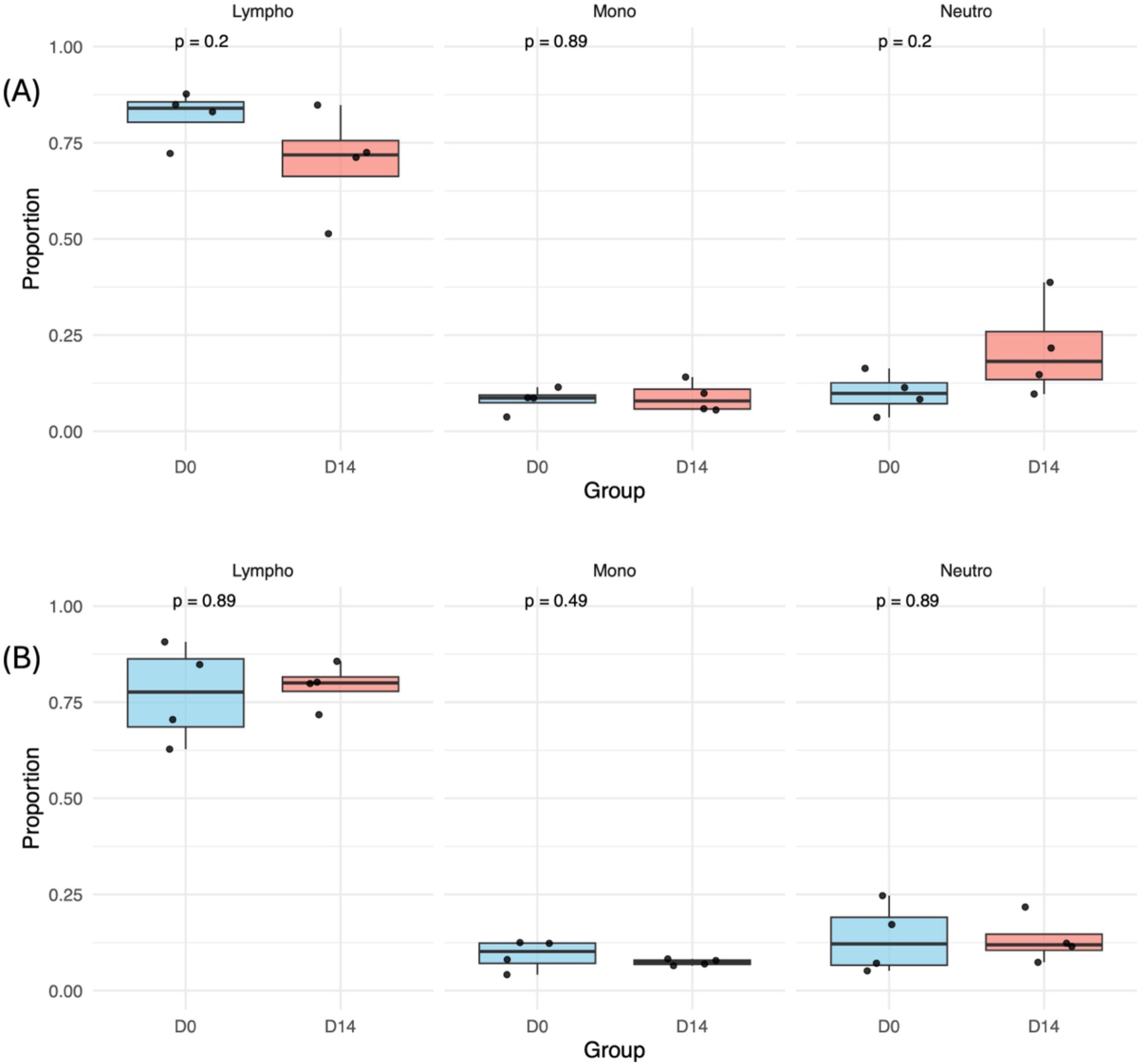
Immune cell-type proportions before and after heat stress. Proportions of immune cell types were estimated at baseline (D0) and after 14 days (D14) under heat stress (HS; panel A) and thermoneutral (TN; panel B) conditions. Estimates were obtained using a reference-based deconvolution approach (EpiDISH) with a bovine immune cell methylation reference panel (t-test, *p* < 0.05). Lympho: Lymphocyte; Mono: Monocyte; Neutro: Neutrophil.

After filtering steps - removing CpGs with insufficient coverage, overlapping known SNPs, and not consistently detected across all libraries (see Methods) - a final set of 1,359,970 common CpGs_10-500_ was retained. These sites were used to assess methylation profile similarity between samples using unsupervised hierarchical clustering (Figure 2). In samples collected under heat stress conditions (Figure 2A), libraries clustered distinctly by collection time point, indicating clear differences in DNA methylation profiles in PBMCs before (D0) and after (D14) the heat challenge. Conversely, in thermoneutral conditions (Figure 2B), clustering by time point was less evident, although some individuals still exhibited time point specific grouping.

**Figure 2.**
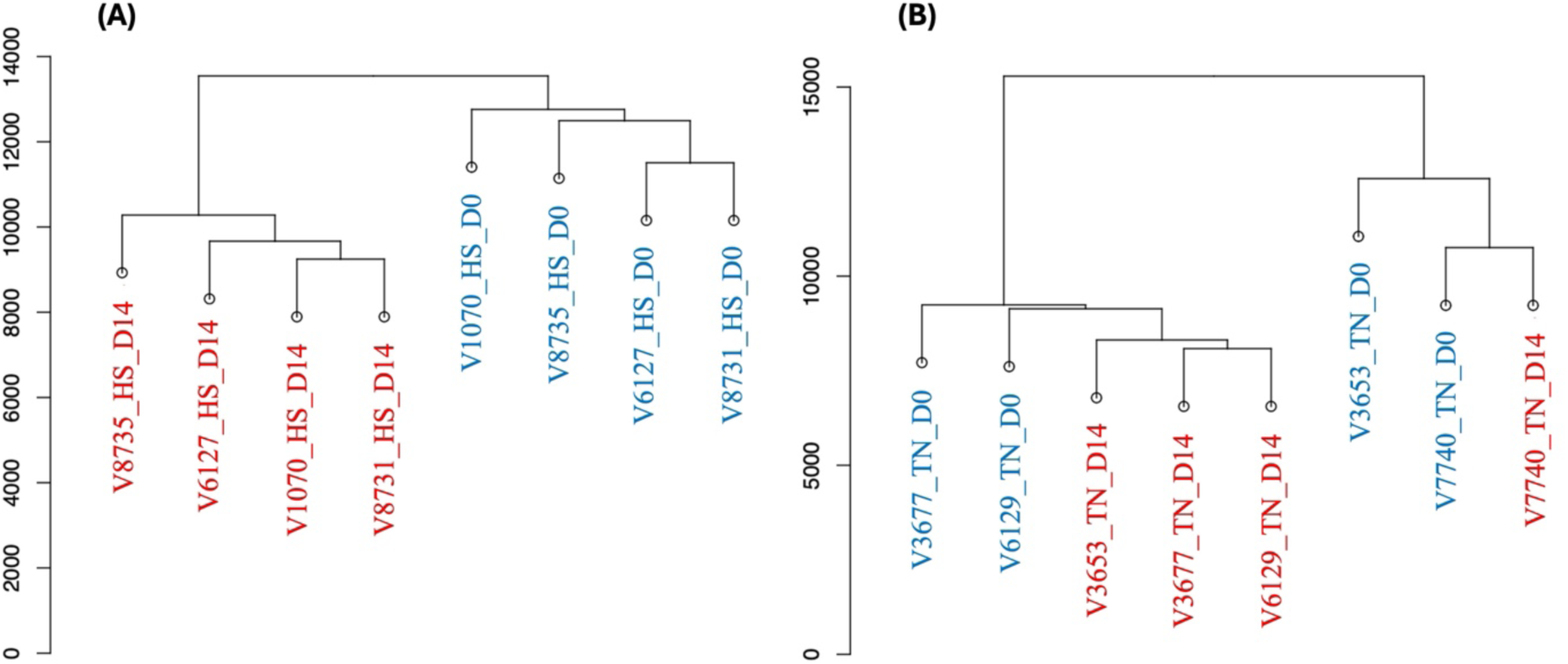
Unsupervised hierarchical clustering based on Euclidean distance calculated from the 1,359,970 common CpGs_10-500_. (A) Clustering of samples collected before (D0, red) and after (D14, blue) the heat stress challenge. (B) Clustering of samples collected at D0 (red) and D14 (blue) from cows maintained under thermoneutral conditions.

### Heat stress induced DNA Methylation Changes: Single-Cytosine Resolution and Genomic Context Analysis

To identify heat stress-induced differentially methylated cytosines (DMCs), DNA methylation profiles in PBMCs were compared between D0 and D14; same cows within each group (Figure 3). Cows exposed to heat stress and those maintained under thermoneutral conditions were analyzed separately using two algorithms (methylKit and DSS), applying a minimum methylation difference of 25% and an adjusted p-value ≤ 1% (see Methods). Importantly, for each cow, the pre-challenge sample (D0) served as an individual baseline, allowing the assessment of changes in DNA methylation in response to heat stress or thermoneutral conditions.

**Figure 3.**
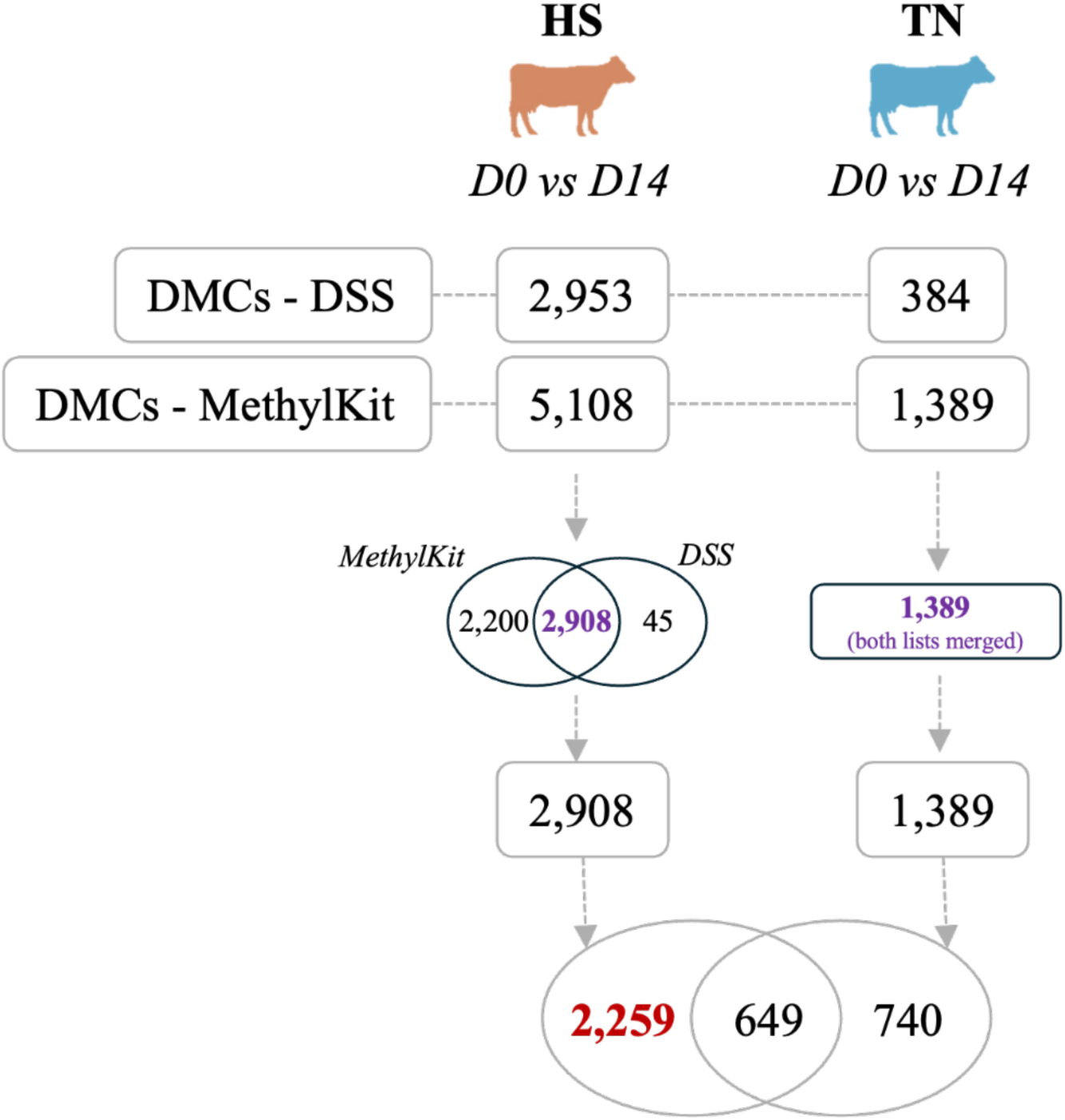
Identification and filtering of differentially methylated cytosines (DMCs).

In the heat stress group (D0 vs. D14), DSS identified 2,953 DMCs, while methylKit detected 5,108 DMCs. The intersection of these two datasets yielded a credible set of 2,908 DMCs (*Figure* 3). In the thermoneutral group, DSS identified 384 DMCs, and methylKit detected 1,389 DMCs. After merging these results, a total of 1,389 unique DMCs were retained (*Figure* 3). The lower number of DMCs identified by both algorithms under thermoneutral conditions indicates a less pronounced methylation change over time and/or feeding status compared to the heat-stressed cows, where a higher number of DMCs were consistently detected by both algorithms.

To enrich for methylation changes specifically induced by heat stress and to exclude methylation variation unrelated to thermal conditions, we filtered the broader heat stress DMC set (2,908 DMCs) by removing all sites that overlapped with those found under thermoneutral conditions (1,389 DMCs) (*Figure* 3). This filtering step resulted in a final set of 2,259 DMCs likely to be associated with the heat stress response, which were subsequently retained for downstream analyses. A detailed characterization of the 2,259 DMCs is provided in Supplementary File S2.

With the final 2,259 DMCs dataset, we observed a pronounced global hypomethylation following heat stress exposure, with 98.23% DMCs showing decreased methylation levels at D14, post heat stress challenge (Figure 4A). Analysis of genomic feature co-localization revealed that 29.3% of DMCs were in intergenic regions, while the remaining 70.7% were found within gene bodies (Figure 4B). Notably, approximately 45.5% of DMCs were associated with intergenic and/or intronic regions - areas often enriched for regulatory elements involved in gene expression control (Jones, 2012; Borsari et al., 2021). Supporting this regulatory potential, the majority of DMCs (84.5%) were located within CpG islands (Figure 4C), which are linked to transcriptional regulation. In contrast, DMCs did not show a strong enrichment for any repeat family (Figure 4D), suggesting that methylation changes were not preferentially associated with repetitive elements.

**Figure 4.**
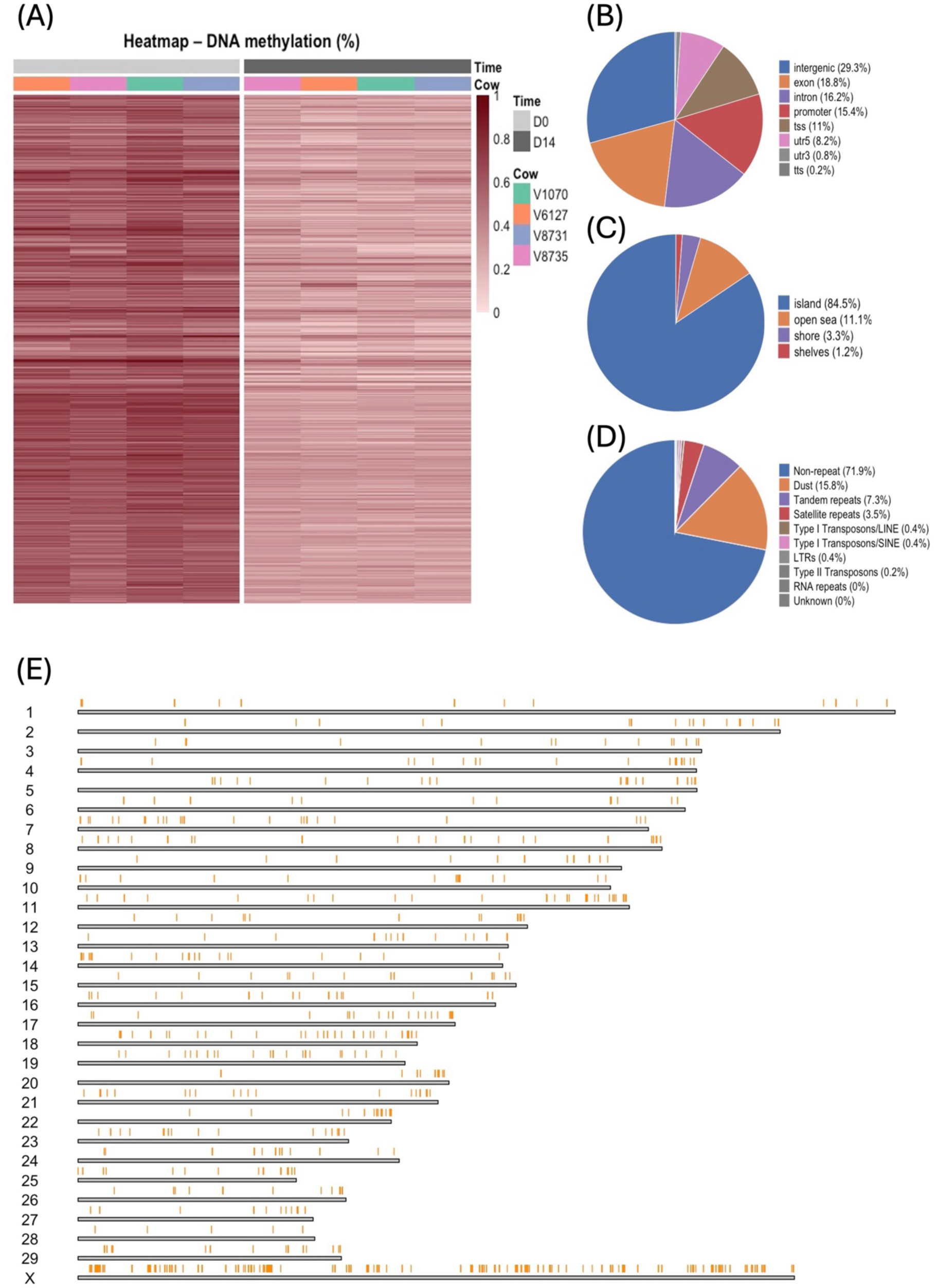
DNA methylation levels and genomic feature enrichment of the 2,259 DMCs. (A) Heatmap showing the percentage of DNA methylation at each DMC for all animals at two time points: D0 (baseline) and D14 (after heat stress challenge). Rows represent CpG sites, and columns represent individual animals. (B–D) Pie charts showing the proportion of DMCs overlapping gene features (B), CpG island classes (C), and repeat types (D). (E) Genome-wide distribution of 2,259 DMCs across bovine chromosomes.

An analysis of the chromosomal distribution of the final set of 2,259 DMCs revealed a slight bias in the terminal regions of several chromosomes (Figure 4E). Notably, we also identified enriched DMC regions on the X chromosome (Figure 4E), which is known to play a key role in the genetic architecture of complex traits in dairy cattle (Sanchez et al., 2023).

### Heat stress induced DNA Methylation Changes: Linking Differentially Methylated Cytosines to Gene Targets

We performed functional enrichment analysis using the list of genes targeted by all common CpGs_10-500_ as the background (18,528 genes) and the list of genes targeted by all DMCs (2,259 DMCs targeting 605 genes) as the main gene set, which revealed 145 significantly enriched Gene Ontology Biological Processes (GOBP) (FDR < 0.05; Supplementary File S3). Although no terms were directly linked to immune system function, most enriched processes were associated with nervous system development, neurogenesis, neuron projection and differentiation, as well as transcriptional regulation, biosynthetic activity, developmental programs, and anatomical structure morphogenesis (Supplementary File S3).

To further refine the search for epigenetic biomarkers and to better understand the impact of heat stress on the immune system of lactating cows, we used the list of genes targeted by all DMCs (n = 605; Supplementary File S2) to identify GO terms associated with the immune system and heat stress response. Genes with no detectable expression in immune cells were filtered out using the Human Protein Atlas (Uhlén et al., 2015) (https://www.proteinatlas.org/), resulting in the prioritization of 27 candidate genes (Table 1).

**Table 1.**
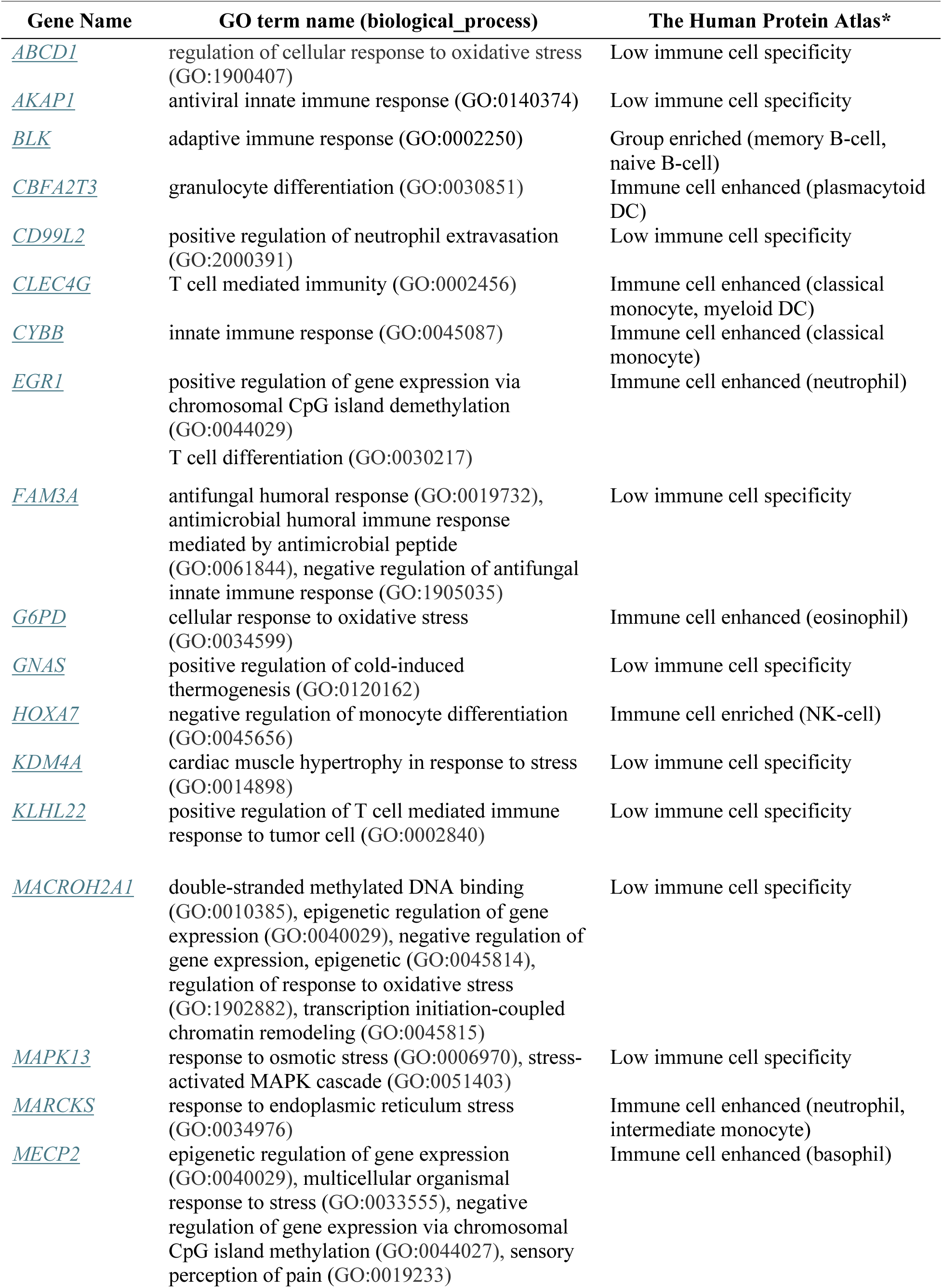
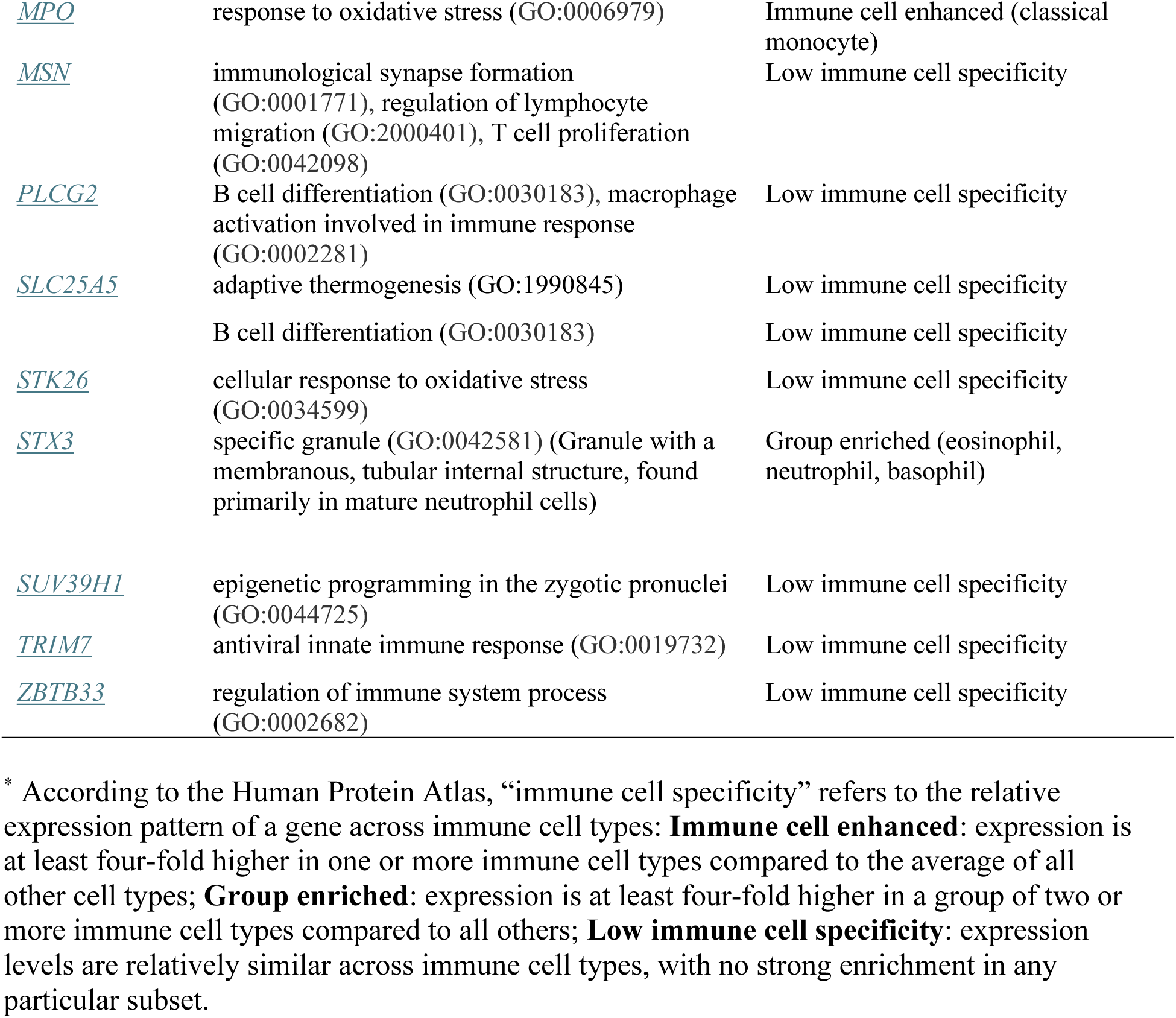
Candidate genes identified through GO term searching, their associated biological processes, and immune cell expression profiles from the Human Protein Atlas (Uhlén et al., <u>2015)</u> (https://www.proteinatlas.org/)

### Heat stress induced DNA Methylation Changes: Linking Epigenetic Signatures to Functional Mechanisms

To identify genomic regions most likely to influence gene regulation under heat stress, we applied a region approach by detecting Differentially Methylated Region (DMR). To do so, we aggregated DMCs into contiguous genomic regions (see Methods for additional details), allowing the detection of coordinated methylation changes that may have greater functional significance than isolated DMCs. Therefore, by prioritizing DMRs in our biomarker analysis, we may enhance our chances of identifying methylation changes that directly affect gene expression and cellular responses to heat stress.

A total of 215 DMRs were identified from the 2,259 DMCs analyzed, all showing a loss of methylation (Supplementary File S4), consistent with the observed global hypomethylation. Analysis of their chromosomal distribution revealed a less pronounced enrichment in the terminal regions of chromosomes, in contrast to what was observed for the DMCs (Figure 4E/DMCs and 5/DMRs). Notably, approximately half of the DMRs are located on the X chromosome (Figure 5).

**Figure 5.**
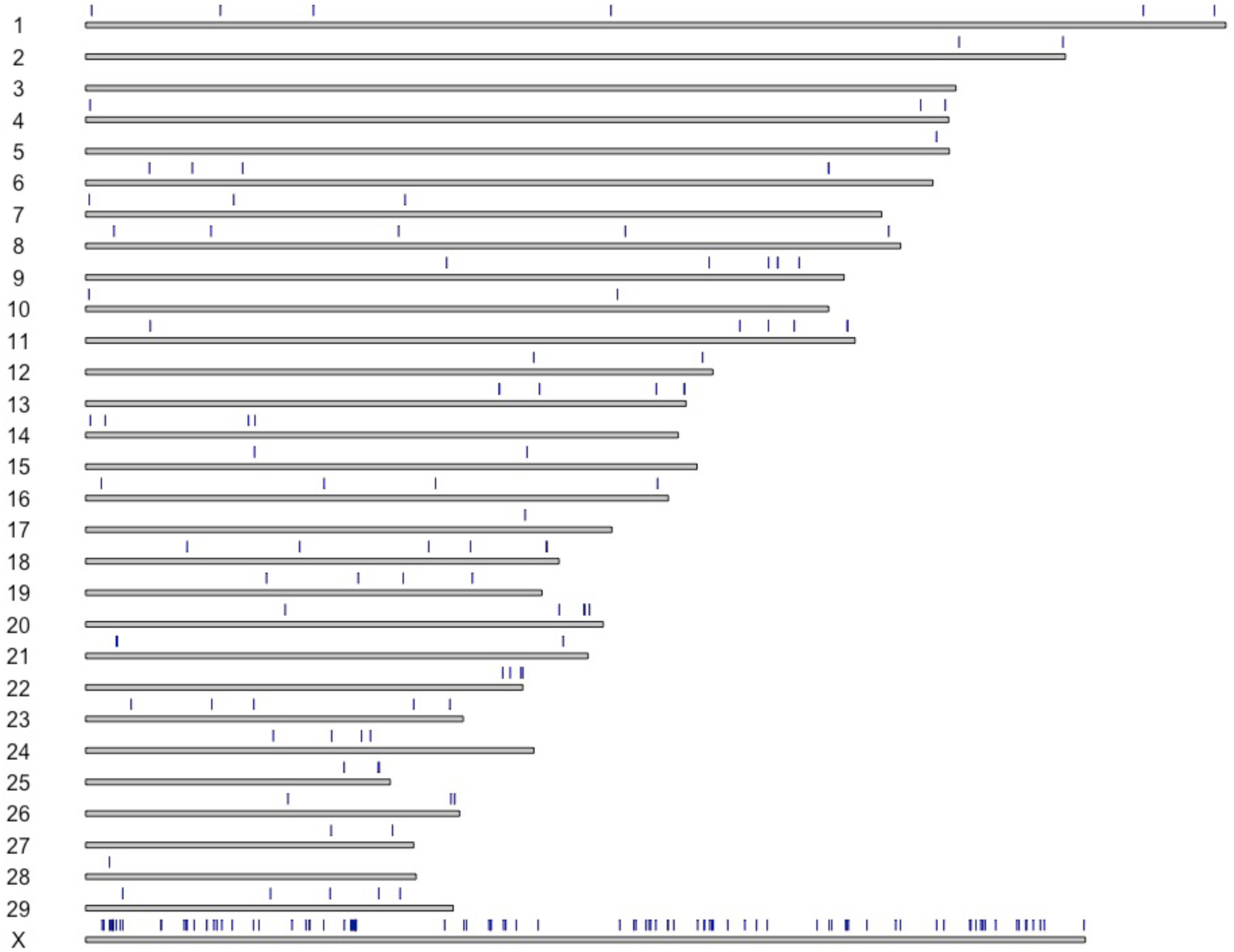
DNA methylation levels and genomic feature enrichment of 215 DMRs. Genome-wide distribution of 215 DMRs across bovine chromosomes.

To better characterize the regions identified toward a better understanding of the functional role of the DMRs identified, we investigated their co-localization with regulatory features (Ensembl Regulatory features – ARS-UCD1.3). A total of 75 DMRs are co-localized with regulatory elements (promoter, open chromatin regions and enhancer) (Supplementary File S4). Interestingly, out of the 75, the majority are co-localized with enhancers (n=45; 60%) followed by promoters (n=26; 34.66%) and open chromatin regions (n=4; 5.33%). Out of the 75 DMRs co-localized with regulatory elements, six encompassed candidate genes highlighted in Table 1 and were further investigated (Figure 6).

**Figure 6.**
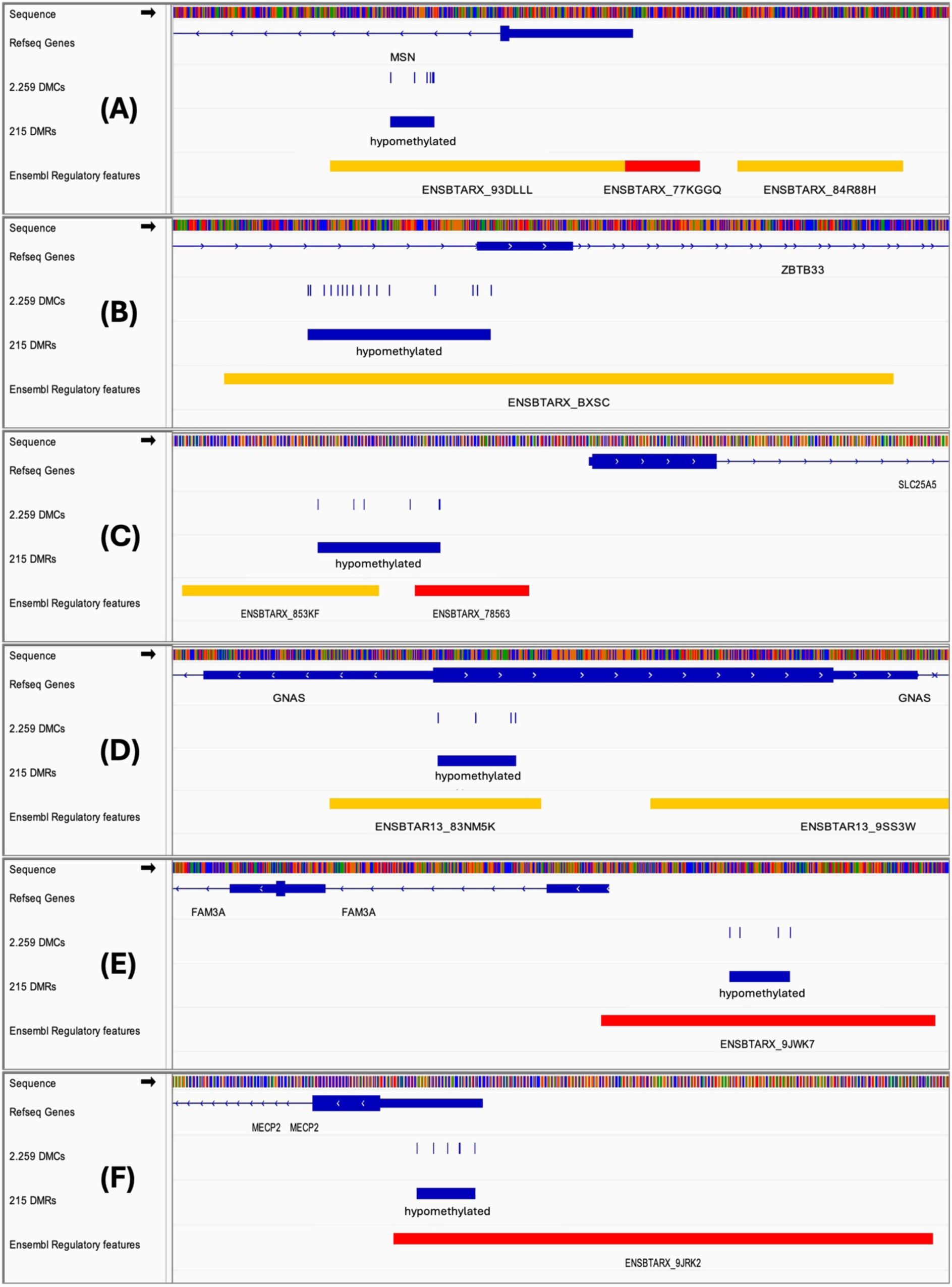
IGV screenshot showing six DMRs co-localized with regulatory elements. (A) Hypomethylated DMR overlapping an enhancer in the *MSN* gene; (B) hypomethylated DMR overlapping an enhancer in the *ZBTB33* gene; (C) hypomethylated DMR overlapping an enhancer and promoter in the *SLC25A5* gene; (D) hypomethylated DMR overlapping an enhancer in the *GNAS* gene; (E) hypomethylated DMR overlapping a promoter in the *FAM3A* gene; (F) hypomethylated DMR overlapping a promoter in the *MECP2* gene. Yellow bars indicate enhancers, and red bars indicate promoters.

## Discussion

A marked impact of heat stress was observed on both physiological and production parameters in dairy cows. While no differences were observed between HS and TN groups at baseline (D0), exposure to HS significantly elevated rectal temperatures and respiratory rates by 4% and 240%, respectively, by D14. Although dry matter intake decreased over time in both groups, milk yield was consistently lower in HS cows, averaging 14% less than TN cows despite similar feed intake. This production decline under HS was accompanied by pronounced metabolic and inflammatory responses, with plasma insulin, lipopolysaccharide-binding protein, tumor necrosis factor alpha, and C-reactive protein increasing by 46–202%, and fecal calprotectin by 27%, highlighting the systemic stress and inflammation induced by heat exposure. Collectively, these findings underscore the detrimental effects of HS on cow performance and metabolic health (Ruiz-Gonzalez et al., 2023).

### Impact of Heat Stress on Epigenetic Regulation of cow’s PBMCs

Our deconvolution analysis of DNA methylation data indicated that heat stress did not lead to statistically significant changes in the proportions of major immune cell types in PBMC preparations (p > 0.05). Although not statistically significant, cows exposed to heat stress showed a tendency toward reduced lymphocyte abundance accompanied by a relative increase in neutrophils (Figure 1A), a pattern that was not observed under thermoneutral conditions, where only a slight decrease in monocytes was detected (Figure 1B). Overall, these results suggest that the epigenetic changes associated with heat stress in PBMCs are unlikely to be driven by shifts in immune cell composition but instead may reflect regulatory responses occurring within immune cells.

Chromosomal distribution of the 2,259 DMCs showed a slight bias in terminal/subtelomeric regions (Figure 4E). In humans, subtelomeric DNA methylation is known to change with age (Bacalini et al., 2021) and has been associated with age-related diseases (Hu et al., 2019). These regions are also rich in repetitive CpG dinucleotides, making them particularly susceptible to epigenetic regulation via DNA methylation (Blasco, 2007; Mikkelsen et al., 2007; Bacalini et al., 2021). Because RRBS targets CpG-dense regions (e.g., promoters and CpG islands), subtelomeric areas can be oversampled when CpG-rich (Nakabayashi et al., 2023). Consistently, previous studies have reported higher methylation levels in CpG islands near telomeres and subtelomeres, which could reflect both biological mechanisms and methodological bias (Zhou et al., 2016). Therefore, the observed bias in terminal/subtelomeric regions should be interpreted with caution, as it may result from a combination of biological effects and methodological artifacts.

When analyzing chromosomal distribution, we also observed an enrichment of DMCs and DMRs on the X chromosome (Figure 4E/DMCs and 5/DMRs). In females, one copy of the X chromosome is inactivated early in development, leading to the silencing of one X chromosome in each cell, which is then clonally maintained in all daughter cells (Lyon, 1961; Plath et al., 2002). Thus, hematopoietic stem cells (HSCs) in the bone marrow, from which lymphocytes, monocytes, and neutrophils are derived, already possess an established inactive X chromosome. Consequently, all immune cells originating from a single HSC share the same inactive X. However, previous studies have shown that, due to the presence of multiple HSC clones with independently inactivated X chromosomes, the immune cell population in adult females can exhibits a mosaic pattern of X-chromosome inactivation (Sharp et al., 2000; Carrel and Willard, 2005).Therefore, although the X chromosome plays a key role in the genetic architecture of complex traits in dairy cattle (Sanchez et al., 2023), these results should be interpreted with caution.

In our study, cow PBMC methylomes were compared over a 14-day interval. To minimize the detection of DMCs attributable to differences in nutrient intake rather than heat stress, cows in the thermoneutral group were pair-fed to match the intake of their heat-stressed counterparts. By excluding all DMCs that were also observed in the thermoneutral (pair-fed) group sampled at the same time points, we increased the chances that the retained DMCs in the credible set could be predominantly associated with heat stress exposure rather than differences in feed intake or temporal effects.

Based on the co-localization of the DMCs detected herein and genomic features, different observations corroborates that they may influence gene expression: (i) a substantial proportion of DMCs were located within gene bodies; (ii) the majority of DMCs resided in CpG islands, which are well known to contribute to transcriptional regulation (Long et al., 2016); and (iii) approximately half of the DMCs were found in intergenic or intronic regions, known to harbor regulatory elements (Jones, 2012; Borsari et al., 2021). In contrast, DMCs did not exhibit strong enrichment within any specific repeat family, suggesting that these methylation changes did not target repetitive sequences and were likely not involved in genome destabilization.

Enrichment analysis of the genes targeted by the 2,259 DMCs (605 genes) revealed overrepresentation of biological processes mainly related to nervous system development, neurogenesis, neuron projection and differentiation, as well as transcriptional regulation, biosynthetic activity, developmental programs, and anatomical structure morphogenesis. Although these functions are classically associated with neural tissues, it is important to highlight that neural signals are known to modulate immune responses and, a considerable number of molecular mediators and signaling pathways are shared between nervous and immune systems (Dantzer, 2017). The enrichment of these pathways in PBMCs therefore suggests a systemic adjustment affecting both immune competence and cellular plasticity. Moreover, the convergence of neural and immune signaling mechanisms may reflect a broader adaptive response to stress, indicative of neuroimmune crosstalk that contributes to the maintenance of homeostasis under adverse thermal conditions.

### Molecular Pathways Underlying Heat Stress Response: Insights from Candidate Genes

The negative impact of heat stress on lactating cow’s immune system is quite well documented (Fabris et al., 2017; Dahl et al., 2020; Gupta et al., 2022), with a known role in both adaptive and innate immune pathways. Herein, we have identified seven candidate genes that highlights this interplay: *BLK*, *PLCG2*, *INAVA*, *CLEC4G*, *MPO*, *CYBB* and *TRIM7* (Table 1). For instance, the *BLK* and *PLCG2* genes, are central regulators of B-cell receptor signaling, suggest that adaptive immune activation may be modulated under heat stress (Xu et al., 2007; Bernal-Quirós et al., 2013). In parallel, genes linked to innate immunity, such as the Innate immunity activator (*INAVA)* gene, which regulates NOD-like receptor signaling a key component of immune response (Chen et al., 2009), and the *CLEC4G* a C-type lectin involved in pathogen recognition and negatively regulation T cell-mediated immunity, and in apoptotic cell clearance by macrophages (Yang et al., 2018), indicate that heat stress may alter immune response mechanisms. The *MPO* and *CYBB* genes, are both essential for reactive oxygen species (ROS) generation in neutrophils (Aratani, 2018; Belambri et al., 2018; Zeng et al., 2019), highlighting the link between immune response and oxidative stress; generally exacerbated during heat exposure. The *TRIM7* gene acts as an inhibitor of apoptosis, playing a role in cell death regulation and viral defense (Fan et al., 2021; Gonzalez-Orozco et al., 2024). The detection of these seven candidate genes that highlights this interplay between both adaptive and innate immune pathway, suggests an impact of heat stress on immune signaling, inflammatory modulation, and oxidative balance, potentially contributing to the immunosuppression observed in heat-stressed dairy cows (Dahl et al., 2020).

Five candidate genes identified are linked to transcriptional regulation and chromatin remodeling, highlighting the role of epigenetic in the response and adaptation to environmental challenges: *EGR1*, *MECP2*, *SUV39H1*, *KDM4A*, and *ZBTB33* (Table 1). The *EGR1* is a transcription factor which can be induced by stress signals (Shajahan-Haq et al., 2017) involved in cell survival, proliferation, and repair (Wu et al., 2009), suggesting that its activation may represent a first line/early regulatory mechanism in the heat stress response. In addition, well known epigenetic modulators such as *MECP2*, *SUV39H1*, *KDM4A*, and *ZBTB33* influence chromatin accessibility and transcriptional plasticity (Blattler et al., 2013; Honer et al., 2024). DNA methylation changes in these regulators could affect the epigenomic landscape, affecting gene expression in the cells under heat stress.

Interestingly, developmental transcription factors including *SOX11*, *HOXA7*, and *PRDM12* were also detected as candidate genes targeted by the DMCs (Table 1). While these genes are traditionally studied in developmental contexts, emerging evidence highlights their continued relevance in adult tissues. For example, the *SOX11* gene, is essential for both embryonic and adult neurogenesis (Wang et al., 2013). Similarly, the *PRDM12* gene, a member of the PRDM family of epigenetic regulators involved in neurogenesis, is selectively expressed in developing nociceptors and remains active in mature nociceptors in adults (Latragna et al., 2019). Together, these findings suggest that these transcription factors (which may retain functional roles beyond development), potentially contributes to adult stress responses.

Three genes related to cytoskeletal organization and cell adhesion, processes that are essential for immune cell function, we also identified: *MSN*, *PKP3*, and *MARCKS* (Table 1). *MARCKS*, *PKP3*, and *MSN* play pivotal roles in regulating the actin cytoskeleton and cell junction dynamics (Brudvig and Weimer, 2015; Fang et al., 2022; Gupta et al., 2023), which are essential for maintaining immune cell morphology and migration. Interestingly, supporting our findings, Hirata et al.(Hirata et al., 2012) reported that *MSN* knockout mice displayed decreases in both T and B cells in the peripheral blood, highlighting the critical role of *MSN* in lymphocyte homeostasis and, consequently, in proper immune function.

Seven genes from the list are involved in metabolic adaptation: *G6PD*, *GNAS*, *SLC25A5*, *ABCD1*, *AKAP1*, *PPARGC1A* and *FAM3A* (Table 1). The *G6PD* gene encodes a key enzyme of the pentose phosphate pathway, is essential for generating NADPH and protecting cells against oxidative stress, which is elevated during thermal exposure (Tchouagué et al., 2019). The other candidates such as *GNAS*, *SLC25A5*, *ABCD1*, *AKAP1*, *PPARGC1A* and *FAM3A*, also plays a role in processes such as mitochondrial activity, ATP transport, energy and lipid metabolism (Chen et al., 2005; Liang and Ward, 2006; Clémençon et al., 2013; Liu et al., 2020; Mallack et al., 2022; Yan et al., 2023), sustaining cellular energy demands; crucial during lactation and heat stress adaptation. Together, these findings indicate that heat stress induces a coordinated modulation of metabolic pathways, likely aimed at maintaining cellular energy balance and antioxidant capacity, thereby buffering immune and physiological functions essential for animal survival and productivity.

It is important to highlight that *GNAS* is an imprinted locus with complex, tissue-specific expression patterns (Kelsey, 2010); imprinting patterns vary across tissues reflecting the locus’s functional diversity. In many tissues, the Gαs transcript is predominantly expressed from the maternal allele, whereas other transcripts, such as XLαs, are mainly expressed from the paternal allele (Bastepe, 2007; Kelsey, 2010; Cui et al., 2021). These features make *GNAS* particularly interesting for studying heat stress–associated methylation, as differential methylation could influence which allele is expressed in a tissue-specific manner, potentially affecting hormone signaling, energy metabolism, and stress adaptation.

In addition to immune and metabolic pathways, we identified genes involved in hormonal, neuroendocrine, and nervous system responses, highlighting the systemic nature of heat stress adaptation. Specifically, *PDX1* regulates multiple aspects of pancreatic β-cell biology, including differentiation, maturation, function, survival, and proliferation (Baumel-Alterzon and Scott, 2022); *RET*, a receptor tyrosine kinase, has been linked to stress response pathways (Myers and Mulligan, 2004); *ADCYAP1* encodes PACAP, a neuropeptide that modulates neuroendocrine and behavioral stress responses (Ebner et al., 2024) and; *ESRRG*, a nuclear receptor widely expressed in the brain, regulates neuroendocrine and behavioral stress responses in rats (Fox et al., 2025), highlighting its potential role in systemic adaptation to heat stress. Together, these genes underscore the tight integration of endocrine, metabolic, and neural mechanisms in heat stress adaptation.

Moreover, genes such as *STX3*, *CBFA2T3*, *CD99L2*, and *MACROH2A1* may indirectly modulate the secretion of cytokines and hormones during stress adaptation. Specifically, *STX3* has been shown to play an essential role in cytokine trafficking in neutrophil granulocytes (Naegelen et al., 2015); *CBFA2T3* functions as a transcriptional corepressor involved in cell differentiation and gene regulation (Steinauer et al., 2020); CD99L2 regulates a distinct step in leukocyte transmigration(Rutledge et al., 2022); and the histone variant MACROH2A1 contributes to chromatin organization and epigenetic control of gene expression (Corujo and Buschbeck, 2018).

Finally, *MAPK13* and *STK26* plays a central role in stress-responsive cellular pathways. *MAPK13* integrates diverse signals to regulate cell proliferation, differentiation, transcription, and development (Cremades-Jimeno et al., 2025); processes that are likely to be modulated under thermal stress. *STK26* (*MST4*), through its role in autophagic activity, is essential for cellular and organismal responses to harmful stimuli, including infections, and contributes to the regulation of immunity and inflammation(Levine et al., 2011). Together, these genes represent key nodes where heat stress may alter epigenetic regulation to modulate cell survival, stress adaptation, and immune function.

### Linking Epigenetic Signatures to Functional Mechanisms

Recent efforts by the FAANG community (https://www.animalgenome.org/community/FAANG/) have substantially advanced the functional annotation of the bovine genome across multiple tissues, enabling more precise identification of regulatory elements. Our findings underscore the importance of integrating such functional annotations to refine the biological interpretation of candidate epigenetic biomarkers. In this context, we focused on six hypomethylated DMRs overlapping regulatory elements located near key candidate genes identified in this study (Figure 6). These DMRs map to genes with central roles in immune regulation, metabolic homeostasis, and chromatin dynamics. As hypomethylation at regulatory elements is generally associated with increased chromatin accessibility and transcriptional activity, our findings suggest enhanced regulatory responsiveness under heat stress. Specifically, hypomethylation was detected in enhancer regions of *MSN*, *ZBTB33*, and *GNAS*, in enhancer/promoter regions of *SLC25A5*, and in promoter regions of *FAM3A* and *MECP2*.

Hypomethylation within an enhancer of *MSN* may promote its expression and influence lymphocyte homeostasis during thermal challenge. Similarly, reduced methylation in a *ZBTB33* enhancer suggests increased activity of this chromatin regulator, which has been associated with inflammatory modulation in murine models (Chaudhary et al., 2013). For *SLC25A5* epigenetic changes at enhancer and promoter regions point to potential upregulation of this mitochondrial ADP/ATP transporter, a key factor in T cell survival and function *(Gicobi et al., 2021; Yosef et al., 2025)*, thereby linking mitochondrial energetics to immune adaptation.

Among the six candidate genes highlighted (Figure 6), *GNAS* stands out as particularly compelling in the context of heat stress induced epigenetic regulation. As previously discussed, *GNAS* is a complex, imprinted locus with tissue-specific expression patterns (Bastepe, 2007; Kelsey, 2010). Moreover, recent studies in mice demonstrate that disruption of Gαs signaling in CD11c⁺ innate immune cells enhance energy expenditure, promotes adipose tissue beiging, and protects against obesity and insulin resistance, underscoring a critical role for *GNAS* in immune cell-mediated metabolic homeostasis (Zeng et al., 2023). Hypomethylation within a GNAS enhancer may reflect enhanced Gαs signaling, potentially supporting metabolic flexibility and systemic energy balance under heat stress.

As previously discussed, *FAM3A* plays a central role in metabolic homeostasis by regulating ATP synthase activity (Yan et al., 2023). Beyond metabolic regulation, *FAM3A* has been shown to protect mitochondrial function during Endoplasmic reticulum (ER) stress-mediated apoptosis in neuronal HT22 cells (Song et al., 2016). The hypomethylation within a promoter region of the *FAM3A* gene (Figure 6E), suggests its upregulation under heat stress conditions. Since ER stress and mitochondrial dysfunction are key modulators of immune cell activation and survival (Bronner et al., 2015; Pereira et al., 2022), these findings suggest that *FAM3A* may similarly contribute to maintaining PBMC function under heat stress.

Finally, in addition to *MECP2* well-established epigenetic regulator functions, recent studies also highlight its importance in immune system regulation. For example, in a mouse model, (Zalosnik et al., 2021) observed that *MECP2* deficiency exacerbates neuroinflammation and promotes autoreactive responses during autoimmune challenges. Additionally, *MECP2* enhances regulatory T cell resilience to inflammation by maintaining Foxp3 expression (Li et al., 2014). In our study, we detected loss of DNA methylation in a *MECP2* promoter region (Figure 6F) under heat stress, suggesting a potential upregulation of *MECP2* in PBMCs, which could epigenetically modulate immune responses in response to heat stress.

Taken together, the consistent localization of hypomethylated DMRs within enhancers and promoters of genes involved in immune regulation, mitochondrial function, and chromatin organization supports the presence of targeted epigenetic remodeling under heat stress. Rather than isolated events, these modifications may affect regulatory regions of genes that integrate metabolic and immune pathways, suggesting coordinated adjustments at the transcriptional level. While functional validation is required to confirm direct effects on gene expression, the convergence of these epigenetic changes highlights biologically coherent mechanisms that may affect immune competence and metabolic balance in response to thermal challenge.

## Conclusion

Our study provides novel insights into the impact of heat stress on the PBMC epigenome of lactating cows, revealing widespread DNA methylation changes predominantly characterized by hypomethylation. For the first time, we observed: (i) a substantial number of CpGs putatively affected by heat stress – relatively to studies conducted using RRBS approach; (ii) significant enrichment of DNA methylation changes in CpG islands, suggesting a pronounced effect on gene expression; and (iii) notable enrichment of DMCs/DMRs on the X chromosome, uncovering an uncharacterized layer of epigenetic regulation under heat stress conditions. By integrating methylome data with functional annotation, we identified key candidate genes whose epigenetic modulation under heat stress may influence immune, metabolic, neuroendocrine, and stress-response pathways. These alterations likely contribute to the immunosuppression and physiological adaptations reported in heat-stressed cattle, providing a mechanistic link between epigenetic modifications and functional outcomes. Overall, our findings advance the understanding of how heat stress can reshape the dairy cows’ epigenome and highlight candidate epigenetic biomarkers. These results could also inform mitigation strategies, such as targeted nutrient or feed additive supplementation, to prevent DNA demethylation and potentially improve dairy cow welfare under climate-change-associated heat stress.

## NOTES

The Canadian experiment was funded in part by the Agriscience Program (ASP-060) and the CRASD (2019-BL-386). The authors also thank the local “Epipôle” structure for stimulating discussions and valuable scientific exchanges.

